# Biochemical reconstitution of sister chromatid cohesion establishment during DNA replication

**DOI:** 10.1101/2025.02.25.640040

**Authors:** Masashi Minamino, John F. X. Diffley, Frank Uhlmann

## Abstract

Concomitant with DNA replication, the ring-shaped cohesin complex encircles both newly synthesized sister chromatids, enabling their faithful segregation during cell divisions. Our molecular understanding of how cohesin co-entraps both replication products remains incomplete. Here, we reconstitute sister chromatid cohesion establishment using purified budding yeast proteins. Cohesin rings, initially loaded onto template DNA, remain DNA-bound during DNA synthesis and encircle both replication products. Additionally, DNA replication instigates new cohesin recruitment, as a second pathway that generates sister chromatid cohesion. In both scenarios, cohesin often embraces only one of the two replication products, suggestive of a two-step sister chromatid capture mechanism. Sister chromatid co-entrapment occurs independently of replication fork-associated cohesion establishment factors, suggesting a role for the latter during chromatin replication or in facilitating the subsequent cohesin acetylation reaction. Our results make sister chromatid cohesion establishment amenable to direct experimental exploration.

## INTRODUCTION

Concurrently with genome replication during S phase of the eukaryotic cell cycle, the chromosomal cohesin complex establishes physical links between the two replication products^1–4^. These links, known as sister chromatid cohesion, provide the counterforce to the mitotic spindle when chromosomes align on the cell equator, in preparation for their segregation into daughter cells during cell divisions. Cohesin is a ring-shaped protein complex of the Structural Maintenance of Chromosomes (SMC) family^5^. Cohesin rings are topologically loaded onto DNA already before the onset of DNA replication, during G1 phase, with help of a Scc2-Scc4 cohesin loader complex^6–8^. Following DNA replication, individual cohesin rings embrace both sister chromatids^9–11^. What happens during DNA replication, when the replisome moves along chromosomes and encounters cohesin, remains incompletely understood.

Different scenarios have been proposed for how cohesin transitions from encircling one DNA to embracing two sister DNAs at this time. These possibilities are not mutually exclusive, and they include i) replisome passage through cohesin rings^12,13^, ii) transient unloading and then reloading of cohesin complexes behind the fork^13^, iii) *de novo* cohesin loading at the replication fork^7,10^, and iv) cohesin being pushed to replication termination sites for cohesion establishment^14^.

In addition to the cohesin complex, a series of replisome components contribute to sister chromatid cohesion establishment, known as ‘cohesion establishment factors’. They include the Tof1-Csm3-Mrc1 replisome progression complex^15–17^, the structural replisome component Ctf4 with its binding partner, the Chl1 helicase^18–20^, as well as the PCNA sliding clamp and its Ctf18-RFC loader complex^18,21,22^. Complex genetic relationships between these cohesion establishment factors^7,23,24^ suggest that more than one reaction contribute to efficient sister chromatid cohesion establishment.

Additional to entrapping both replication products, cohesin must be acetylated on two conserved lysine residues to ensure that sister chromatid co-entrapment results in enduring sister chromatid cohesion^25,26^. Acetylation stops dynamic cohesin loading and unloading cycles that otherwise characterize cohesin’s chromatin interactions^27–30^, a requirement for stable sister chromatid linkages. Again, the above mentioned cohesion establishment factors contribute in a complex overlapping pattern to the acetylation reaction^24,31^, which is furthermore dependent on transient DNA structures that form during Okazaki fragment maturation^32^.

Biochemical reconstitution is a powerful approach to dissect and understand complex molecular events. Complete eukaryotic DNA replication has been reconstituted using purified components from budding yeast^33,34^, as has the topological loading of cohesin rings onto DNA^6,35^. Combining *in vitro* cohesin loading with DNA replication opens an opportunity to study the establishment of sister chromatid cohesion. A recent report on such reconstituted replisome-cohesin encounters uncovered a replication-coupled cohesin loading reaction, mediated by Ctf4-Chl1^36^. Whether such loaded cohesins topologically entrapped both replication products remained untested. Replisome-cohesin encounters were also studied in *Xenopus* cell free egg extracts, where reported outcomes included replisome passage or cohesin pushing events^3,14,37^. The complex nature of the extracts complicated the molecular dissection of cohesion establishment events, and the characterization of the resultant sister chromatid linkages.

Here, we reconstitute sister chromatid cohesion establishment during DNA replication using purified budding yeast components. Cohesin that is loaded onto DNA before the onset of DNA replication topologically embraces both replication products. Additionally, the replisome promotes cohesion establishment by *de novo* cohesin recruitment during ongoing DNA replication. In both cases, we find that cohesin often entraps only one of the two replication products, consistent with a two-step sister chromatid co-entrapment mechanism. *In vitro* co-entrapment of both replication products occurs independently of replisome-associated cohesion establishment factors, suggesting that the latter play roles during *in vivo* chromatin replication, or are destined to promoting the cohesin acetylation reaction.

## RESULTS

### Cohesin remains DNA bound during DNA replication

We set out to reconstitute sister chromatid cohesion establishment during DNA replication. We began by purifying all budding yeast components required for origin-dependent DNA replication^33,34^. In our experiments, MCM helicase double hexamers were assembled onto an *ARS1* replication origin containing, 3.2 kb circular plasmid DNA template. Next, MCM double hexamers were activated by Dbf4-dependent kinase (DDK). Finally, origin firing factors, DNA polymerases, and accessory proteins were added to initiate DNA replication. [α-^32^P]dCTP was included amongst the nucleotides to facilitate visualization of replication products. Following deproteination, the reactions were then resolved by native agarose gel electrophoresis, and replication products visualized by autoradiography, while total DNA was stained with the DNA dye SYBR Gold. Topoisomerase II (Top II) was included in these reactions to decatenate fully replicated sister DNA circles. This setup resulted in the generation of completely replicated circular DNA products (Figure S1).

We next added cohesin to this replication assay. *In vivo*, cohesin is topologically loaded onto chromosomes before the onset of DNA replication^4,7^. We therefore started by loading cohesin onto our template plasmid DNA, with help of the cohesin loader and ATP, before MCM loading and activation. Cohesin immunoprecipitation at this stage, before initiating DNA synthesis, confirmed that cohesin was successfully loaded onto the template DNA (Figure 1A, sample **a**). We next initiated DNA replication using this cohesin-bound template by adding the remaining replication proteins, which included the cohesion establishment factors Ctf4, Tof1-Csm3 and Mrc1. We additionally included purified Chl1, Ctf18-RFC, as well as Pds5 and Eco1^32^. Following DNA replication, cohesin immunoprecipitation revealed that cohesin was associated with circular replication products (Figure 1A, sample **b**). This observation left unresolved whether cohesin remained DNA-bound during DNA replication, or whether new cohesin was loaded during or following DNA synthesis. To distinguish between these two possibilities, we prepared sample **c**, in which no cohesin was loaded onto the DNA template, but cohesin was present together with its loader during the DNA replication incubation. In this case, cohesin barely associated with replication products. The likely reason is the presence of 250 mM potassium glutamate in the replication buffer, high ionic strength conditions that disfavor cohesin loading (Figure 1A, sample **c**). Therefore, cohesin-bound to replication products in sample **b** must have derived from cohesin that was loaded onto DNA at the beginning of the experiment. The fraction of cohesin-bound DNA before DNA replication (stained with SYBR Gold) was comparable to the fraction of cohesin-bound replication products (visualized by autoradiography), suggesting that there was little cohesin loss during DNA replication. These observations suggest that cohesin that encircles DNA before DNA replication remains DNA bound during DNA replication. This conclusion was also recently reached by Murayama et al. (2024)^36^.

**Figure 1.**
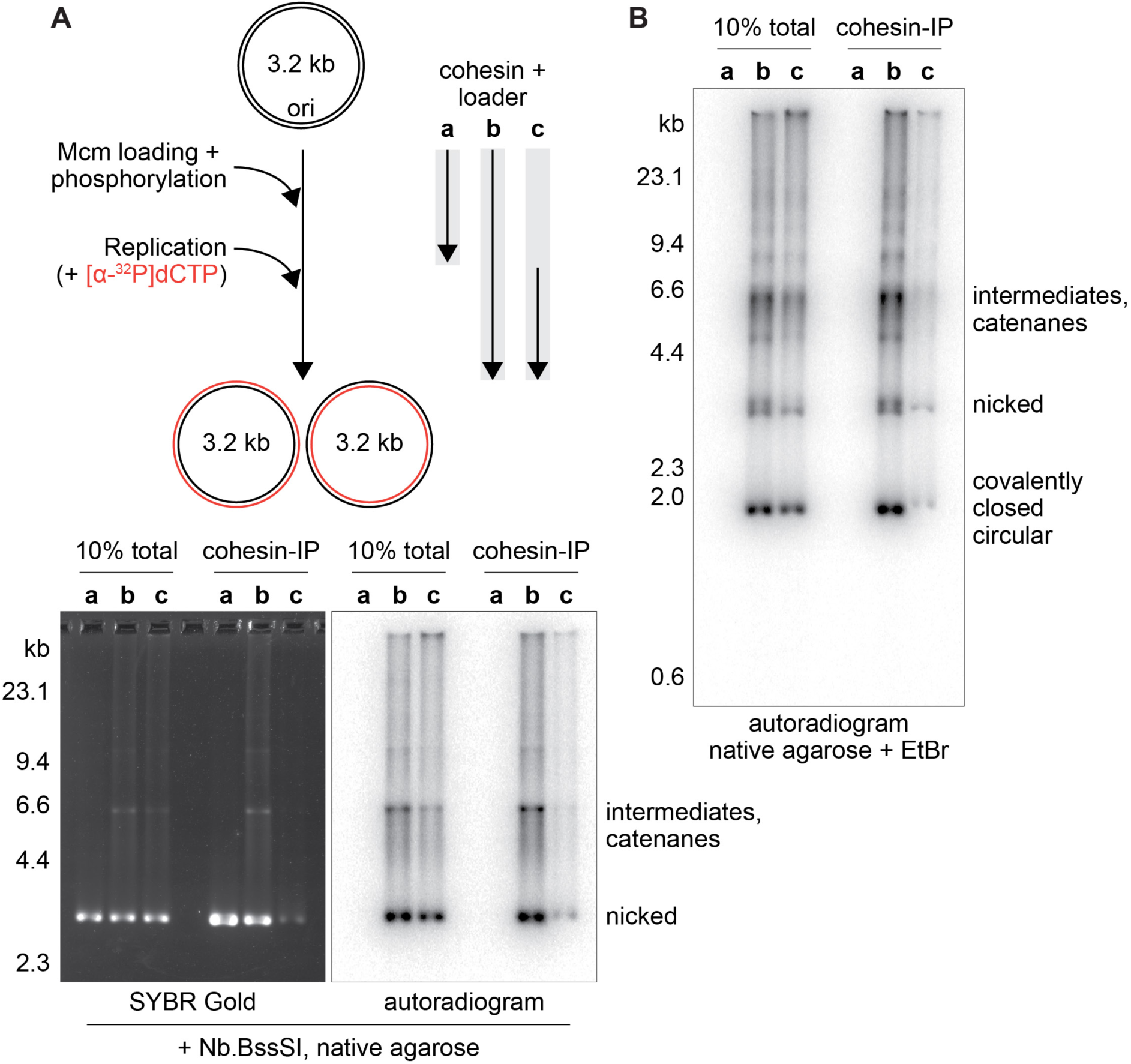
Cohesin remains DNA bound during DNA replication. (A) Schematic of the experiment in which cohesin is **a** loaded onto unreplicated DNA during origin licensing, **b** loaded onto unreplicated DNA during origin licensing, followed by DNA replication, **c** added after origin licensing only during the replication reaction. Total products (total) and cohesin-bound DNAs (cohesin-IP) were separated following nicking enzyme Nb.BssSI treatment by agarose gel electrophoresis. Total DNA was visualized by SYBR Gold staining, replication products by autoradiography of incorporated [α-^32^P]dCTP. See also Figure S1 for further characterization of the DNA replication reaction. (B) The experiment was repeated but DNA samples were separated, without nicking, on an ethidium bromide (EtBr)-containing agarose gel. Covalently closed circular DNAs migrate faster due to EtBr-induced supercoiling, confirming that cohesin remains bound to fully replicated sister DNAs. See also Figure S2 for characterization of the fully replicated species, as well as an experiment to probe the contribution of cohesion establishment factors to cohesin retention during DNA replication.

### Cohesin associates with completely replicated DNA circles

The above experiment showed that cohesin remains bound to *in vitro* replicated and decatenated DNA circles. While our template DNA is supercoiled, topoisomerases present in the replication reaction cause its partial relaxation. To facilitate the comparison between input and recovered DNA, irrespective of their supercoiling state, we had treated all samples in Figure 1A with a nicking enzyme that converts all DNAs into relaxed circles before agarose gel electrophoresis. To investigate whether cohesin remains associated with completely replicated covalently closed replication products, we omitted nicking enzyme treatment and instead resolved the replication products on an ethidium bromide containing agarose gel. Ethidium bromide intercalates and thereby leads to supercoiling only of covalently closed circular DNAs (Figure S2A). As seen in Figure 1B, our replication reaction proceeded to yield such covalently closed circular DNA replication products, and these were bound by cohesin. This observation demonstrates that cohesin that is loaded onto DNA before DNA replication remains bound to DNA throughout all DNA synthesis, ligation and decatenation steps that lead to two completely replicated sister DNA products.

We investigated whether cohesin retention during DNA synthesis requires any of the replisome-associated cohesion establishment factors. However, omitting Ctf4-Chl1, Tof1-Csm3, or Mrc1 from the replication reaction did not noticeably alter the efficiency with which cohesin was retained on replication products (Figure S2B). Taken together, the results so far suggest that cohesin remains DNA bound during the process of complete *in vitro* plasmid DNA replication.

### Biochemical reconstitution of sister chromatid cohesion establishment

An important unanswered question from the above experiments is whether, following DNA replication, cohesin entrapped one or both replication products. Size analysis of the replication products, required to answer this question, is made difficult by the many DNA binding proteins contained in the replication reaction. To circumvent this problem, we introduced a cohesin variant into our experiments that can be covalently circularized using cysteine-specific crosslinking (6C cohesin)^7^. 3 engineered cysteine pairs allow covalent closure of the three cohesin ring interfaces using the bi-cysteine crosslinker BMOE, while a control 5C variant lacks one of these cysteines. We confirmed that 6C cohesin could be efficiently crosslinked, and that crosslinking following topological loading resulted in SDS-resistant retention of 6C cohesin, but not 5C cohesin, on DNA (Figure S3A-C). The SDS resistant nature of 6C cohesin’s DNA entrapment allowed us to use this detergent to remove all other proteins from the replication products and analyze their cohesion status by gel electrophoresis.

We loaded 6C cohesin onto the 3.2 kb plasmid and used it as the template for DNA replication. Following replication, we added BMOE to covalently circularize cohesin. Then we denatured all proteins by incubation with 1% SDS. Following denaturation, we diluted the sample to reduce the SDS concentration and enable cohesin immunoprecipitation. In the first iteration of this experiment, we then eluted cohesin-bound replication products from the antibody beads using proteinase K (Figure 2A). Separation of these eluates by agarose gel electrophoresis showed retention of replication products by 6C cohesin treated with BMOE, but not by 6C cohesin if the crosslinker was omitted, or by 5C cohesin. This result confirms that cohesin topologically embraces *in vitro* synthesized DNA replication products.

**Figure 2.**
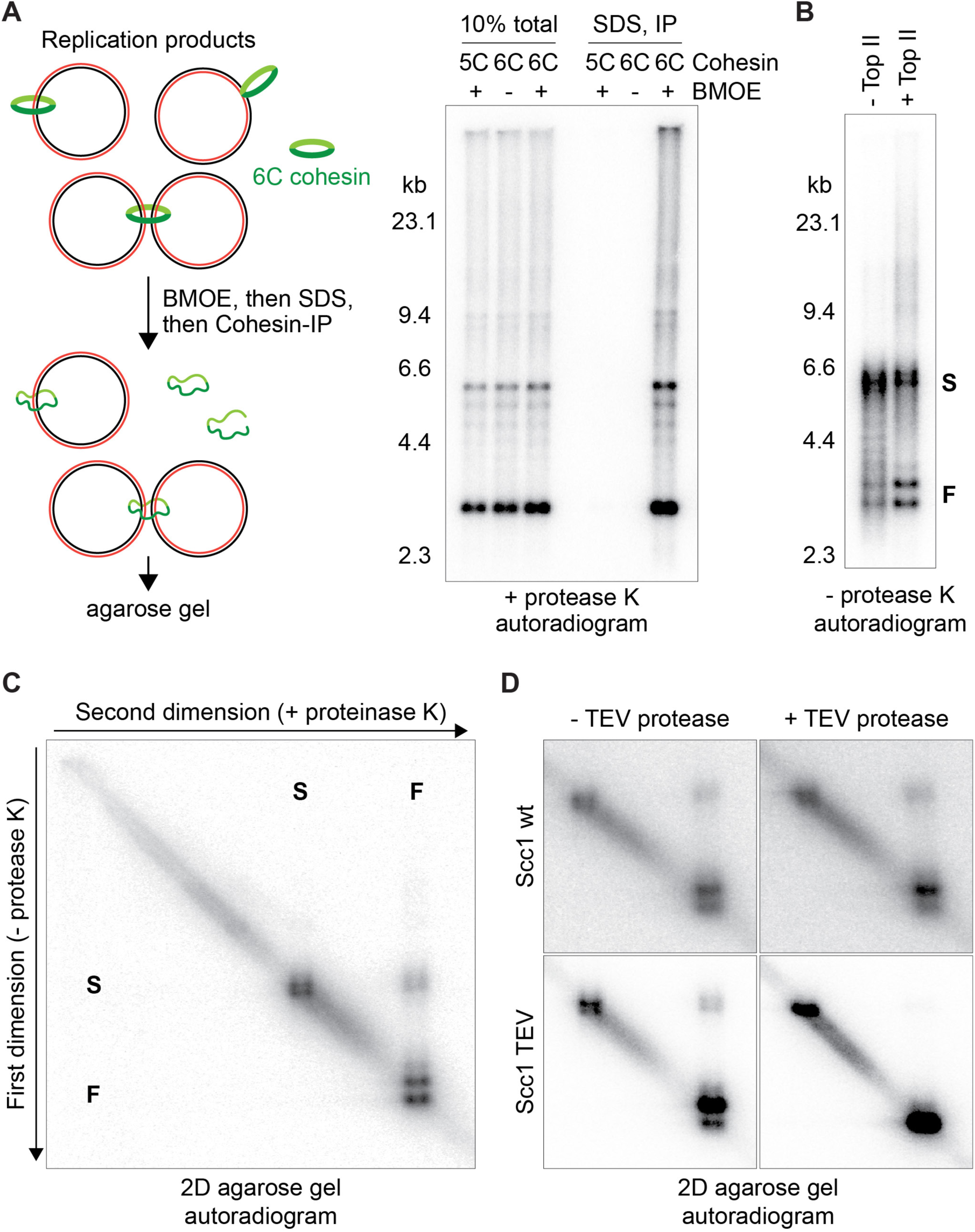
Biochemical reconstitution of sister chromatid cohesion establishment. (A) Schematic of replication products bound by 6C cohesin, and the consequences of its crosslinking and denaturation. Autoradiogram of total replication products formed from 5C or 6C cohesin-bound template DNA (total), and of cohesin immunoprecipitation (IP) following SDS denaturation. (B) Replication product sizes following 6C cohesin crosslinking. DNA replication was performed in the presence or absence of Top II. Fast migrating products (**F**) of approximately the size of the 3.2 kb plasmid, as well as slow migrating products (**S**) of around double the size are indicated (The Pif1 helicase was added to this reaction to facilitate replication termination without Top II^45^). (C) Two-dimensional (2D) gel electrophoresis following replication of a 6C cohesin-bound template DNA, crosslinking, denaturation, and cohesin immunoprecipitation. (D) as (C), but 6C cohesin without or with an Scc1 TEV protease recognition site was used, and samples were treated without or with TEV protease, before 2D gel electrophoresis. See also Figure S3 for further characterization of the 6C cohesin crosslinking experiment.

We next repeated the above experiment, this time eluting cohesin-bound DNA replication products by merely increasing the SDS concentration, but without proteinase K addition. If sister chromatid cohesion was established during DNA replication, the denatured but covalently closed cohesin rings should connect both replication products. Agarose gel electrophoresis of this eluate showed two prominent groups of products. Faster migrating products close to where we expect monomer 3.2 kb circles (Figure 2B, **F** products), and slower migrating products at approximately double that size (**S** products). To create an expectation for where two conjoined replication products would migrate, we performed a replication reaction from which we omitted Top II. Without Top II, products around the size of **S** products increased in intensity, at the expense of **F** products, consistent with the interpretation that the fast and slow migrating bands correspond to monomer and interlinked dimer replication products.

To address whether the **S** band contained pairs of replication products that were held together by cohesin rings, rather than being catenanes, we separated 6C cohesin-associated replication products, generated in the presence of Top II, by two-dimensional (2D) gel electrophoresis. The first dimension was conducted as before, separating the fast and slow migrating populations on a native agarose gel. This gel lane was excised and applied to a perpendicular gel where the products migrated through a proteinase K containing stripe before being separated along an SDS-containing agarose gel (Figure 2C). Any protein-dependent sister chromatid cohesion should be destroyed by proteinase K before separation in the second dimension. While most of the replication products showed comparable migration in the first and second dimensions, forming a diagonal, a distinct fraction of the **S** products were converted to **F** products following proteinase K treatment. This observation suggests that part of the 6C cohesin-associated replication products were protein-dependent dimers, i.e. they were most likely held together by the 6C cohesin ring.

To test whether protein-dependent plasmid dimers were indeed held together by cohesin rings, we inserted a TEV protease recognition sequence into the Scc1 subunit of the 6C cohesin ring (Scc1 TEV, Figure S3D)^38^. We then repeated the replication-coupled cohesion establishment experiment using 6C cohesin containing either wild type Scc1, or Scc1 TEV. Following crosslinking and denaturation, we treated the cohesin immunoprecipitates with or without TEV protease before analyzing the samples by 2D gel electrophoresis (Figure 2D). Protein-dependent dimers were no longer observed following TEV protease treatment of Scc1 TEV-associated replication products, but persisted in mock treated samples, or if TEV protease was used to treat replication products bound to wild type Scc1. This experiment confirms that *in vitro* replication of a cohesin-bound DNA template results in replication product dimers that are topologically held together by the cohesin ring.

### The specificity of *in vitro* sister chromatid cohesion establishment

Next, we needed to establish whether the two DNAs held together by cohesin rings were indeed the two sister chromatids produced at a replication fork, rather than two unrelated DNAs that cohesin might have embraced. Therefore, we asked whether both DNAs in cohesin-linked dimers were replication products, or whether one of the two might be an unreplicated template DNA circle that is present in the replication reaction in excess. To investigate this possibility, we took advantage of the Dam methylation status of the template DNA isolated from *E. coli*. Methylation of both DNA strands makes the template a target for the Dpn I restriction endonuclease. Following *in vitro* replication, the two sister chromatids are each hemi-methylated and thereby turn resistant to Dpn I treatment. We confirmed that our template DNA, containing 16 Dpn I recognition sites, was efficiently degraded following Dpn I treatment (Figure 3A). On the other hand, similar Dpn I treatment of the dimer replication products, associated with 6C cohesin, showed that these were resistant to Dpn I treatment. This observation shows that cohesin-dependent DNA dimers consist of two newly synthesized replication products.

**Figure 3.**
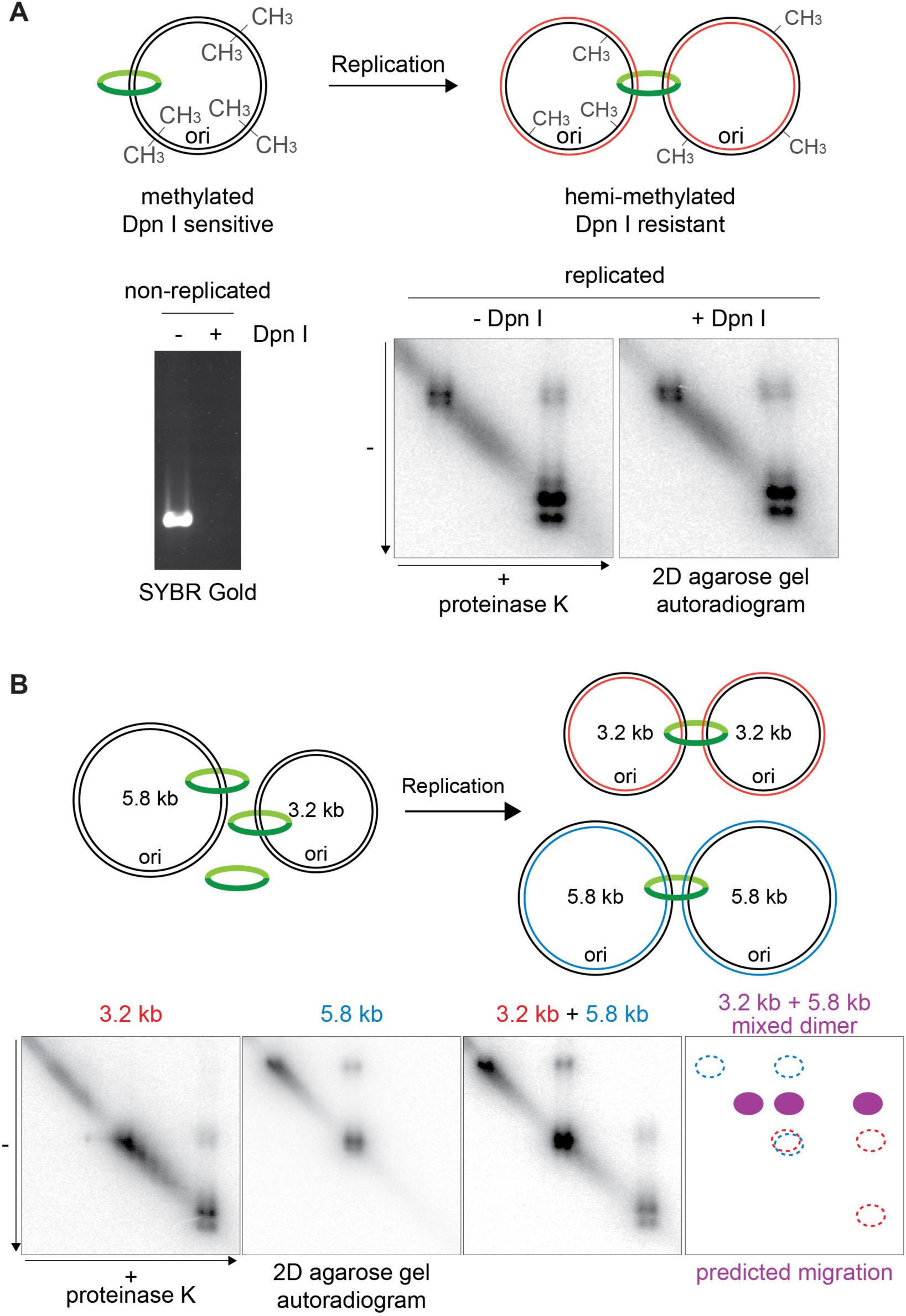
The specificity of sister chromatid cohesion establishment. (A) Replication schematic of a methylated DNA template, and experiments to test the Dpn I restriction enzyme sensitivity of the template DNA, as well as of 6C cohesin-associated replication products analyzed by 2D gel electrophoresis. (B) Schematic of a replication reaction containing two template DNAs of different sizes, and an experiment to analyze 6C cohesin-associated replication products of reactions containing either, or both, templates using 2D gel electrophoresis. The expected band positions produced by mixed dimers, i.e. migrating between the two homodimer sizes in the first dimension, and dimers resolving into both monomer sizes in the second dimension, are indicated in a schematic. See also Figure S4 for one-dimensional agarose gel electrophoresis of replication products produced in the presence of either, or both, templates.

Cohesin must establish interactions between the two sister chromatids that emanate from a replication fork, rather than between any other newly replicated sequences. To test the specificity of *in vitro* sister chromatid cohesion establishment, we performed a replication reaction that contained two DNA circles of different sizes. We loaded 6C cohesin onto the 3.2 kb, as well as onto a 5.8 kb replication origin-containing plasmid. We then mixed the two templates before MCM loading, activation, and DNA replication. Both templates in this mixed reaction were replicated with comparable efficiencies (Figure S4). We then separated 6C cohesin-associated replication products by 2D gel electrophoresis. This analysis revealed two types of protein-mediated DNA dimers. The two types corresponded to those observed in reactions that contained either only the 3.2 kb or only the 5.8 kb template (Figure 3B). We did not detect a signal at positions where a mixed 3.2 – 5.8 kb dimer would be expected to resolve. From these observations, we conclude that cohesin establishes topological interactions specifically between the two sister replication products that are produced during DNA replication.

### Cohesin often embraces one of the two replication products

In addition to cohesin-mediated DNA dimers, changing from slow to fast migration during 2D gel electrophoresis, 6C cohesin recovered several types of replication products that did not change migration following proteinase K or TEV protease treatment (Figure 2C,D). Namely, proteolysis only partly resolved the slower migrating **S** band towards fast **F** migration, suggesting that part of the dimer replication products were connected by protein-independent links, most likely by catenation. Catenanes form as the product of replication termination and they are partially protected from decatenation if the same two DNAs are also held together by cohesin^39,40^. It is therefore possible that the diagonal **S** band products are DNA dimers that were connected by both cohesin as well as DNA catenation.

Notably, immunoprecipitation of 6C cohesin following DNA replication also retrieved a substantial fraction of monomer circles, migrating in the **F** position in both dimensions during 2D gel electrophoresis. How might monomer replication products be topologically entrapped by cohesin? Sister chromatid cohesion, established between replication products, might be unstable and might be lost during the replication incubation, leaving cohesin embracing only one of the sisters at the time of crosslinking. To investigate this possibility, we performed a replication timecourse experiment. DNA replication was largely complete after 30 minutes of incubation (Figure S5A). We then continued the incubation for an additional 30 and 60 minutes before crosslinking 6C cohesin and retrieving cohesin-bound replication products. 2D gel electrophoresis showed that the fraction of protein-mediated dimers remained unchanged over the course this incubation (Figure S5B). This analysis suggests that cohesin-mediated DNA dimers, once formed, are stable for extended periods. Cohesin-bound monomer products must have therefore already arisen during DNA replication. These products reveal that cohesin does not in all cases entrap both sister chromatids. Frequently, *in vitro* DNA replication results in cohesin entrapping only one of the two replication products.

### Cohesion establishment factors and *in vitro* sister chromatid cohesion establishment

We next explored the contribution of the known cohesion establishment factors to *in vitro* sister chromatid cohesion establishment. To do so, we performed a side-by-side comparison of replicating a 6C cohesin-bound plasmid template in the presence or absence of Ctf4-Chl1, Mrc1, Ctf18-RFC, Eco1 and Pds5. This comparison revealed no difference in the generation of protein-mediated dimer products, as detected by 2D gel electrophoresis (Figure 4A). This observation suggests that *in vitro* cohesion establishment proceeds independently of replisome-associated cohesion establishment factors. The role of these factors therefore pertains either to cohesion establishment in a more complex *in vivo* setting where cohesion establishment must be coordinated with histone inheritance^41^, or their role might be restricted to promoting the cohesin acetylation reaction in which they are also known to be involved^24,31^. We were unable to test the role of Tof1-Csm3 in *in vitro* cohesion establishment, as DNA replication was too inefficient in their absence^34^. Though we note that the contribution of Tof1-Csm3 to cohesion establishment is thought to overlap that of Ctf4-Chl1^23^.

**Figure 4.**
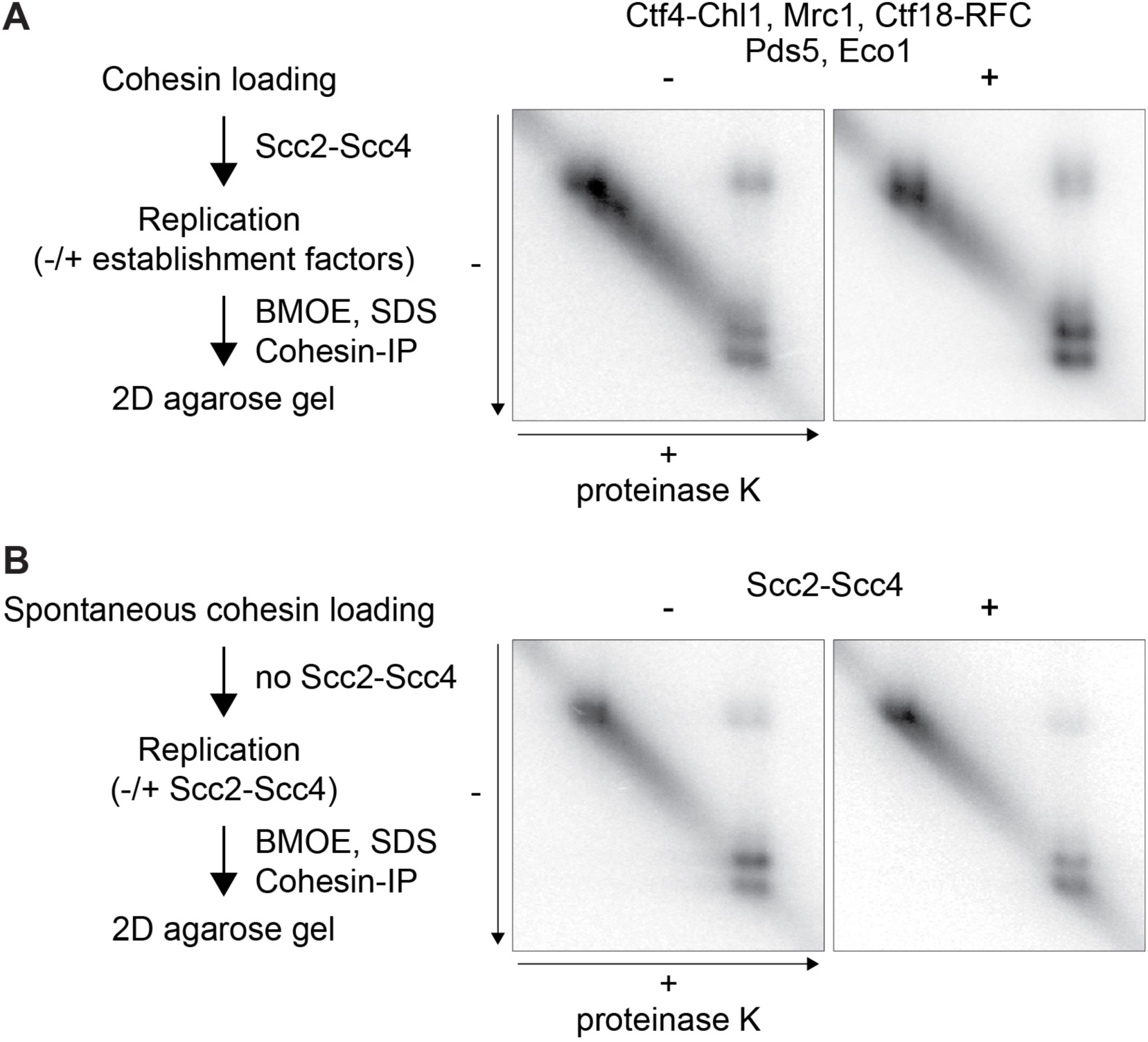
Contribution of cohesion establishment factors and cohesin loader to sister chromatid cohesion establishment. (A) 6C cohesin-bound templates were replicated in the presence or absence of the indicated cohesion establishment factors. Cohesion establishment was assessed following crosslinking, denaturation and cohesin immunoprecipitation by 2D gel electrophoresis. (B) 6C cohesin was loaded onto the template DNA in a reaction without the Scc2-Scc4 cohesin loader, then replication was conducted in the presence or absence of the cohesin loader, and cohesion establishment was monitored as in (A). See also Figure S5 for a timecourse analysis to probe the stability of *in vitro* established sister chromatid cohesion.

In addition to promoting cohesin loading onto chromosomes, the Scc2-Scc4 cohesin loader also contributes to cohesion establishment during S phase^10,32,42^. To investigate a possible role during *in vitro* cohesion establishment, we performed an experiment in which we loaded 6C cohesin onto the template DNA without Scc2-Scc4. *In vitro* cohesin loading proceeds, albeit at a reduced rate, in the absence of the cohesin loader^6,35^. We therefore increased the cohesin loading time, then used the DNA template with loader-independently loaded 6C cohesin in a DNA replication reaction. DNA replication was then performed in either the presence or absence of the cohesin loader. 2D gel electrophoresis of cohesin-bound replication products showed a comparable level of cohesion establishment with or without the cohesin loader (Figure 4B). Taken together, these observations reveal that cohesion establishment during *in vitro* DNA replication occurs independently of cohesion establishment factors and of the cohesin loader.

### Replication-coupled cohesin loading

Our results so far demonstrate cohesion establishment by cohesin that was bound to DNA before DNA replication. We next asked whether soluble cohesin complexes that are present during DNA replication also contribute to sister chromatid cohesion establishment. In Figure 1A, we saw little evidence for cohesin recruitment during DNA replication, at least if replication was carried out at relatively high ionic strength, conditions that impede *in vitro* cohesin loading^6,35^. We therefore repeated DNA replication reactions at a lower salt concentration of 100 mM potassium glutamate, using a cohesin-free template DNA for MCM loading and activation. Cohesin and its loader were then added together with origin firing and replication proteins (Figure 5A). After the replication incubation, we assessed cohesin loading onto DNA replication products. Quantification of total and cohesin-bound DNAs revealed that almost 50% of replication products were bound by cohesin (Figure 5B). This level of DNA recovery was substantially higher than that of template DNA that remained unreplicated in the same incubation, of which only approximately 10% was bound by cohesin. Cohesin loading onto replicated DNA strictly depended on the cohesin loader. These findings suggest that DNA replication prompts cohesin association with replication products to a level that goes well beyond cohesin’s natural tendency to load onto non-replicating DNA.

**Figure 5.**
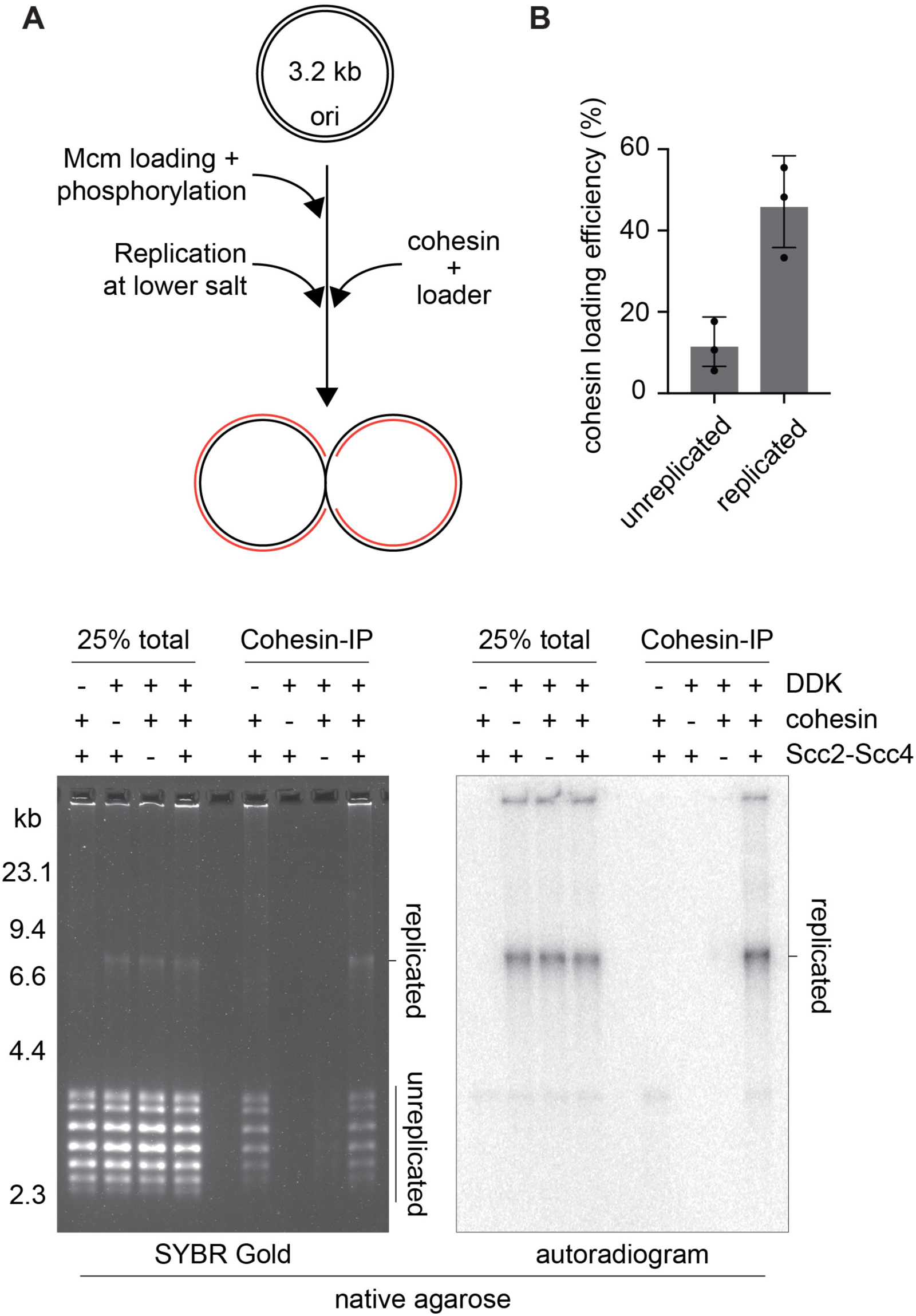
Replication-coupled cohesin loading. (A) Schematic of a replication reaction during which cohesin and the cohesin loader were added at the time of DNA replication. (B) Example gel electrophoretic analysis of the total DNA following the replication incubation, and of the cohesin-bound DNA recovered following cohesin immunoprecipitation (IP). DNA was visualized by SYBR Gold, replication products by [α-^32^P]dCTP autoradiography. Top II was omitted from this experiment, so that late replication intermediates accumulate that migrate distinctly slower than the template DNA. Cohesin loading onto unreplicated and replicated DNA was quantified in three independent biological repeat experiments (black circles). The means (bars) and standard deviations (error bars) are indicated. See also Figure S6 for a characterization of the role of cohesion establishment factors in replication coupled cohesin loading.

### Cohesion establishment by replication-coupled cohesin loading

We lastly wanted to know whether replication-coupled cohesin loading resulted in the establishment of sister chromatid cohesion. To answer this question, we again utilized 6C cohesin and added it together with the cohesin loader during DNA replication. Following completion of DNA replication, we added BMOE to covalently close cohesin rings, performed SDS treatment, and immunopurified topologically entrapped replication products. Analysis by 2D gel electrophoresis revealed the generation of protein-dependent plasmid dimers, suggesting that replication-coupled cohesin loading indeed resulted in the establishment of sister chromatid cohesion (Figure 6A). The dimer species, but not the input template, was resistant to Dpn I treatment, confirming that they consisted of two replication products. Therefore, sister chromatid cohesion can be established by cohesin that is recruited to DNA during ongoing DNA replication.

**Figure 6.**
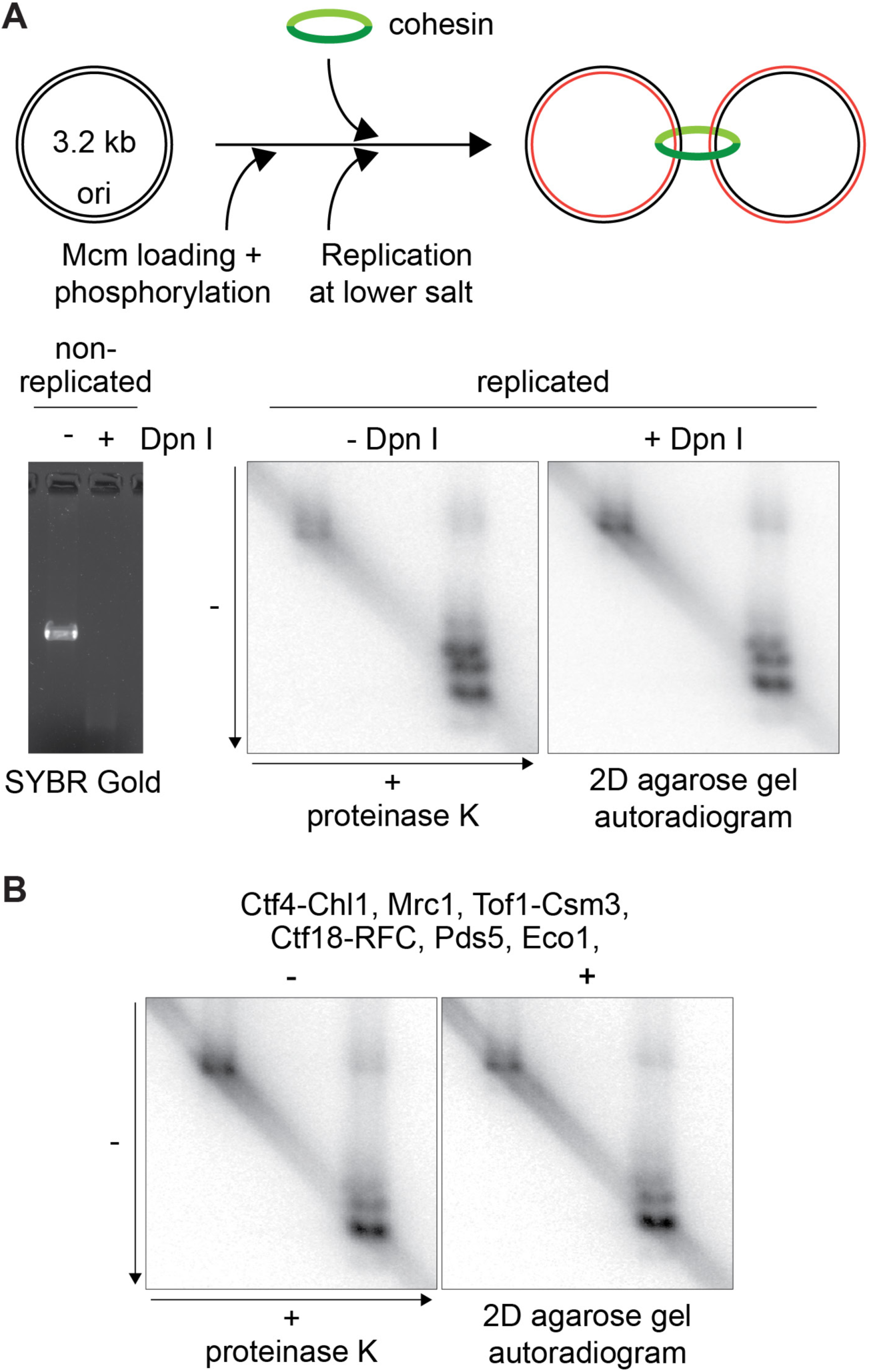
Cohesion establishment during replication-coupled cohesin loading. (A) Schematic of a complete replication reaction, including Pol δ, during which 6C cohesin and the cohesin loader were added at the time of DNA replication. The Dpn I sensitivity of the template DNA was analyzed, as well as of the replication products that were analyzed following cohesin crosslinking and denaturation by 2D gel electrophoresis. (B) As in (A), but DNA replication was conducted in the presence or absence of the indicated cohesion establishment factors.

In addition to the plasmid dimers, 6C cohesin also topologically associated with monomer circular replication products. As was the case during cohesion establishment using pre-loaded cohesin, this observation suggests that a frequent outcome of cohesin loading during DNA replication is cohesin entrapment of only one of the two replication products.

We lastly tested whether cohesion establishment by replication-coupled cohesin loading depended on replication fork associated cohesion establishment factors. The degree of replication-coupled cohesin loading remained unchanged in reaction lacking Ctf4-Chl1, PCNA, Mrc1 or Tof1-Csm3 (Figure S6). Similarly, the level of protein-dependent dimer products formed during DNA replication was similar in reactions containing or lacking Ctf4-Chl1, Mrc1, Tof1-Csm3, Ctf18-RFC, Pds5 and Eco1 (Figure 6B). Sister chromatid cohesion establishment by co-entrapment of the two DNA replication products is therefore a process that occurs independently of replisome-associated cohesion establishment factors.

A recent study reported a role for the Chl1 helicase in promoting replication-coupled cohesin loading^36^. Despite of our efforts, we did not observe a similar Chl1 contribution. The role of Chl1 and its cohesin interactions during cohesion establishment^20^ therefore remains to be further explored.

## DISCUSSION

Here we report the *in vitro* reconstitution of sister chromatid cohesion establishment during DNA replication. Biochemical reconstitution remains a crucial test of our understanding of cellular events. While genetic approaches have identified the many components of the sister chromatid cohesion machinery, we have now confirmed that we know of all the essential parts that are required to ensure that replicated genome copies, synthesized during S phase, remain connected to one another until the time of their segregation in anaphase. At the same time, our biochemical investigations have begun to reveal unexpected insight into the molecular events that lead to sister chromatid cohesion establishment.

### Replication encounters with pre-loaded cohesin

Cohesin that is topologically loaded onto DNA before the onset of DNA replication, results in topological entrapment of the two *in vitro* replicated sister chromatids. Cohesion establishment is equally efficient if replication is performed by a stripped-down replisome, encompassing the bare essentials to carry out DNA synthesis, and lacking many of the components that have been identified as *in vivo* sister chromatid cohesion establishment factors. This observation reveals an intimate relationship between DNA replication and cohesion establishment. As soon as the ring-like architecture of cohesin became known, the possibility emerged that replisomes might simply pass through these rings to leave the replication products trapped inside^12,13^. Recent single molecule observation of real time replisome-cohesin encounters have indeed depicted the replisome apparently slipping through cohesin rings to establish sister chromatid cohesion^43^. It is possible that the same occurs in our bulk biochemical experiments, explaining one of the pathways by which cohesion is established when replication meets cohesin.

In addition to dimer replication products we found that, following DNA replication, cohesin often entrapped only one of the two replication products. While entrapped monomer replication products were a prominent outcome of our reactions, it is not immediately obvious how cohesin might transition from entrapping the template DNA to entrapping only one of the two replication products. Cohesin would have to come off the DNA and then capture only one of the two replication products when getting back on. However, the incidence of single capture events was unchanged in reactions without the cohesin loader, suggesting that cohesin unloading and reloading was not part of the transfer mechanism. An alternative explanation for monomer entrapment is that the cohesin ring ruptured during replisome passage, then closed around only one of the two sisters. The weakest cohesin ring interface at the hinge is known to break at forces in the range of those produced by advancing replisomes^11^. Are single capture products therefore an unavoidable outcome of replisome encounters, with only a fraction of cohesins surviving intact to reach productive sister chromatid cohesion? Or could single capture outcomes be rescued and mature into double capture events? A hint for the latter possibility comes from *in vivo* analyses of how the cohesin loader contributes to sister chromatid cohesion establishment^10,13,42^. Like in our *in vitro* reactions, a basal level of sister chromatid cohesion is achieved without need for the cohesin loader. At the same time, cohesion turns much more robust if the cohesin loader is present at the time of DNA replication. Whether the cohesin loader acts *in vivo* to convert single to double capture events, and why this reaction might have been poorly recapitulated in our *in vitro* setting, remains to be explored.

### Replication-coupled cohesin loading and cohesion establishment

Additional reason to consider single to double capture transitions as a mechanism for cohesion establishment comes from observations with cohesin that is newly recruited to replication products during DNA replication. In this setting, cohesin is again seen embracing only one of the sister chromatids or co-entrapping both. This observation is easiest explained by a sequential DNA-DNA capture mechanism, with a second capture event that is not directly linked to the first. The reaction could be akin to previous *in vitro* observations in which cohesin sequentially captured a double stranded DNA first, followed by a second single stranded DNA^10^. This substrate geometry is reflected at a replication fork where the dsDNA leading strand products lies juxtaposed to the unwound ssDNA lagging strand template. In the future, it will be interesting to explore whether the single capture events are specific to one of the two strands, the leading or the lagging strand. This information should provide clues as to how a sequential capture mechanism might operate at replication forks.

Replication-coupled *de novo* cohesin loading has previously been hypothesized to be one of two means to establish sister chromatid cohesion, based on the contribution of Mrc1 and Ctf18-RFC^7^. However, we find that Mrc1 and Ctf18-RFC are not in fact required for replication-coupled *in vitro* cohesin loading and cohesion establishment. How Mrc1, Ctf18-RFC and the other cohesion establishment factors contribute to building sister chromatid cohesion remains to be understood. *In vivo*, the replisome must coordinate cohesin transfer and DNA capture reactions with histone transfer and redeposition^41,44^. Both reactions involve replisome contacts and require accessible DNA, and several cohesion establishment factors also participate in histone inheritance. Alternatively, the main role of cohesion establishment factors might lie in linking sister chromatid co-entrapment to the cohesin acetylation reaction^24,32^.

Our *in vitro* studies are beginning to reveal how cohesin intersects with the molecular events that take place during DNA replication. In the future, it will be important to develop methods that can follow cohesin behavior during replisome encounters in an *in vivo* setting. These studies will then allow us to complete our molecular understanding of the chromosome replication and segregation cycle.

## ACKNOWLEDGMENTS

We would like to thank M. Douglas and J. Hill for unpublished reagents, the Crick Fermentation Science Technology Platform, as well as our laboratory members for discussion and critical reading of the manuscript. This project received funding through Wellcome Trust Investigator Awards (219527/Z/19/Z to JFXD and 220244/Z/20/Z to FU), and The Francis Crick Institute, which receives its core funding from Cancer Research UK, the UK Medical Research Council, and the Wellcome Trust (cc2002 to JFXD and cc2137 to FU). M.M. was supported by an EMBO Long Term Fellowship and by the Japan Society for the Promotion of Science.

## AUTHOR CONTRIBUTIONS

M.M., J.F.X.D. and F.U. conceived the study. M.M. performed all the experiments, M.M., J.F.X.D. and F.U. wrote the manuscript.

## KEY RESOURCES TABLE

**Table 1.**
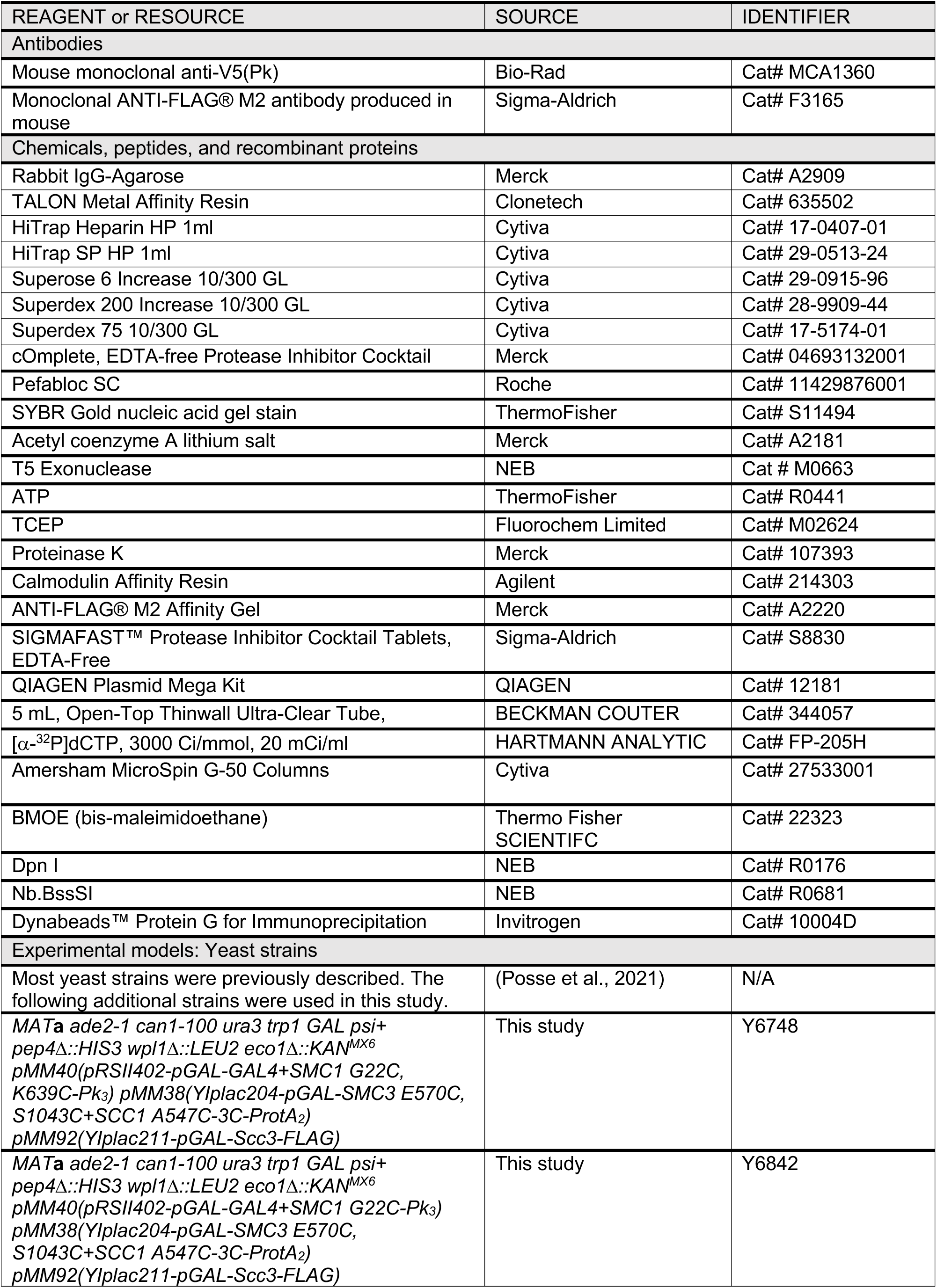
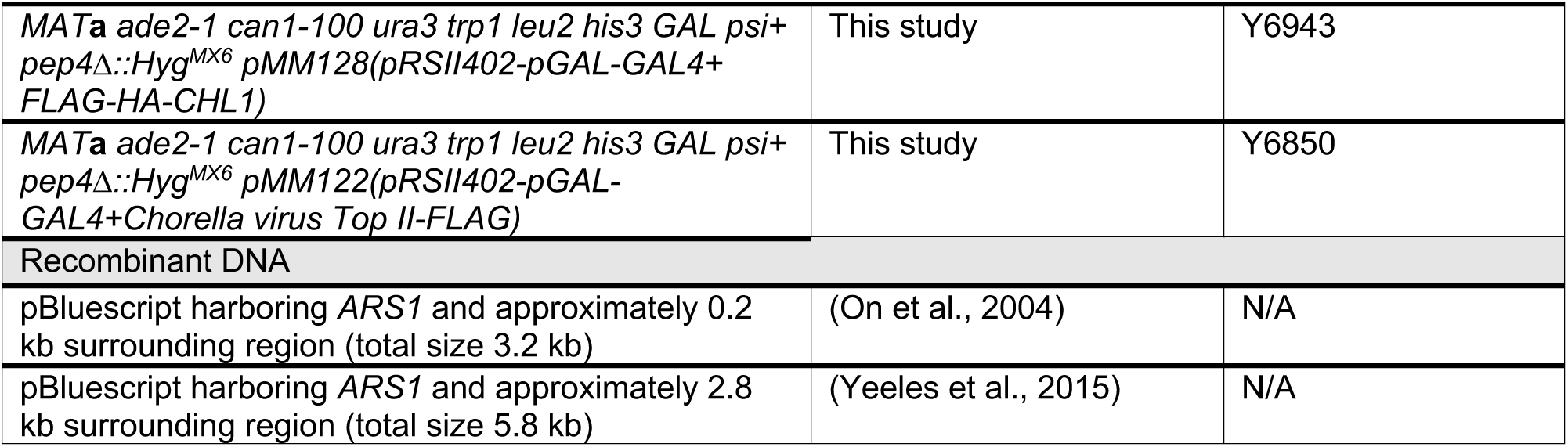

## EXPERIMENTAL MODEL AND SUBJECT DETAILS

### Yeast strains

Budding yeast *S. cerevisiae* strains used in this study are listed in Key Resources Table. Cells were grown in YP medium containing 2% glucose (YPD), 2% raffinose, or 2% raffinose + 2% galactose as carbon source at 30 °C. The conditions used for induction of protein expression are detailed under the respective subsections of the Methods details.

## METHOD DETAILS

### Protein expression and purification

Budding yeast replication proteins (ORC, Cdt1-Mcm2-7, Cdc6, DDK, S-CDK, RPA, Dpb11, Cdc45, GINS, Top I, Top II, Ctf4, Pol ε, Mcm10, Tof1-Csm3, Mrc1, RFC, PCNA, Pol δ, Fen1, Cdc9, Pol α, Sld2, Sld3-Sld7)^46^, Pif1 and Dna2^47^, Eco1, Ctf18-RFC, Pds5^32^, and cohesin and the Scc2-Scc4 cohesin loader complexes^35^ were purified following published protocols.

#### 6C Cohesin purification

Budding yeast cells overexpressing Smc1(G22C, K639C)-Pk, Smc3(E570C, S1043C), Scc1(A547C)-Protein A, and Scc3-FLAG were grown in YP medium containing 2% raffinose as the carbon source to an optical density of 1.0 at 30 °C. 2% galactose was then added to the culture to induce protein expression, and cells were further grown for 2 hours. Cells were collected by centrifugation, washed with deionized water, and suspended in cohesin buffer (50 mM HEPES-KOH pH 7.5, 20% glycerol) containing 300 mM NaCl, 0.5 mM TCEP, 2 mM MgCl_2_, 0.5 mM Pefabloc, as well as cOmplete-EDTA protease inhibitor cocktail. The cell suspension was frozen in liquid nitrogen, then cells were broken in a cryogenic freezer mill. The cell powder was thawed on ice, and further cohesin buffer containing 300 mM NaCl, 0.5 mM TCEP, 0.5 mM MgCl_2_ and protease inhibitors was added. The lysate was clarified by centrifugation at 20,000 x g for 1 hour. RNase A and benzonase were added to a final concentration of 0.3 μg/ml and 1.25 U/ml, respectively, to the clarified lysate. The lysates were transferred to pre-equilibrated Rabbit IgG agarose resin and incubated for 2 hours. The resin was washed with cohesin buffer containing 300 mM NaCl, 0.5mM TCEP, and 2 mM MgCl_2_, then incubated in cohesin buffer containing 300 mM NaCl, 0.5 mM TCEP, 10 mM MgCl_2_ and 1 mM ATP for 15 minutes. The resin was washed again with cohesin buffer containing 300 mM NaCl, 0.5 mM TCEP, and 2 mM MgCl_2_, then incubated overnight in the same buffer containing 10 μg/ml PreScission protease. The eluate was collected, and loaded onto a HiTrap Heparin column, equilibrated with cohesin buffer containing 300 mM NaCl and 2.5 mM TCEP. The column was developed with a linear gradient from 300 mM to 1 M NaCl in cohesin buffer containing 2.5 mM TCEP. The peak fractions were pooled and loaded onto a Superose 6 Increase 10/300 GL gel filtration column that was equilibrated and developed with cohesin 6C gel filtration buffer (20 mM Tris-HCl pH 7.5, 150 mM NaCl, 10% Glycerol, 2.5 mM TCEP). The peak fractions were combined and concentrated by ultrafiltration.

#### 5C and 6C cohesin TEV purification

Cysteine pairs to covalently close the cohesin ring were introduced as described^48^. Budding yeast cells overexpressing Smc1(G22C, K639C)-Pk or Smc1(G22C)-Pk, Smc3(E570C, S1043C), Scc1(A547C)-CBP harboring a TEV protease site (ENLYFQG) in place of the second separase site (SVEQGRR), and Scc3-FLAG were grown in YP medium containing 2% raffinose as the carbon source to an optical density of 1.0 at 30 °C. 2% galactose was then added to the culture to induce protein expression, and cells were further grown for 2 hours. Cells were collected by centrifugation, washed with deionized water, and suspended in cohesin buffer containing 300 mM NaCl, 0.5 mM TCEP, 2 mM MgCl_2_, 0.5 mM Pefabloc, as well as cOmplete-EDTA protease inhibitor cocktail. The cell suspension was frozen in liquid nitrogen, then cells were broken in a cryogenic freezer mill. The cell powder was thawed on ice, and further cohesin buffer containing 300 mM NaCl, 0.5 mM TCEP, 0.5 mM MgCl_2_ and protease inhibitors was added. The lysate was clarified by centrifugation at 20,000 x g for 1 hour. CaCl_2_, RNase A, and benzonase were added to final concentrations of 2 mM, 0.3 μg/ml, and 1.25 U/ml, respectively. The lysate was transferred to pre-equilibrated CBP affinity resin and incubated for 2 hours. The resin was washed with cohesin buffer containing 300 mM NaCl, 0.5 mM TCEP, 2 mM CaCl_2_ and 2 mM MgCl_2_, then incubated in cohesin buffer containing 300 mM NaCl, 0.5 mM TCEP, 2 mM CaCl_2_, 10 mM MgCl_2_ and 1 mM ATP for 15 minutes. The resin was washed again with cohesin buffer containing 300 mM NaCl, 0.5 mM TCEP, 2 mM CaCl_2_, and 2 mM MgCl_2_, then incubated in cohesin buffer containing 300mM NaCl, 0.5 mM TCEP, 2 mM EDTA, and 2 mM EGTA to elute the proteins. The eluate was collected and further purified using HiTrap Heparin and Superose 6 Increase 10/300 GL chromatography, as described above for 6C Cohesin.

#### Chlorella virus Top II purification

During the course of our experiments, we noticed that budding yeast Top II interferes with DNA replication in reactions carried out at lower salt concentrations. As an alternative to yeast Top II, we purified *Chlorella* virus Top II ^49^ as described below. The *Chlorella* virus enzyme proficiently supported DNA replication at low salt concentrations and was used in these experiments.

Budding yeast cells overexpressing *Chlorella* virus Top II were grown in YP medium containing 2% raffinose as the carbon source to an optical density of 1.0 at 30 °C. 2% galactose was then added to the culture to induce protein expression, and cells were further grown for 2 hours. Cells were collected by centrifugation, washed with deionized water, and suspended in CV Top II buffer (25 mM Tris-HCl pH 7.2, 10% glycerol, 400 mM NaCl, 0.01% NP-40, 1 mM DTT) containing 0.5 mM Pefabloc, as well as cOmplete-EDTA protease inhibitor cocktail. The cell suspension was frozen in liquid nitrogen, then cells were broken in a cryogenic freezer mill. The cell powder was thawed on ice, and further CV Top II buffer containing protease inhibitors was added. The lysate was clarified by centrifugation at 20,000 x g for 1 hour. RNase A was added to a final concentration of 0.3 μg/ml to the clarified lysate. The lysate was transferred to pre-equilibrated anti-FLAG M2 affinity gel and incubated for 2 hours. The resin was washed with CV Top II buffer, then incubated in the same buffer containing 10 mM MgCl_2_ and 1 mM ATP for 15 minutes. The resin was washed again with CV Top II buffer, then incubated in the same buffer containing 0.5 mg/ml FLAG peptide. The eluate was loaded onto a Superdex 200 Increase 10/300 GL gel filtration column that was equilibrated and developed with CV Top II gel filtration buffer (25 mM Tris-HCl pH 7.2, 150 mM NaCl, 10% Glycerol, 0.01% NP-40, 1 mM DTT). The peak fractions were combined.

#### Chl1 purification

Codon optimized budding yeast Chl1 was expressed and purified as previously described ^50^. We noticed that increased Chl1 yield due to codon optimization led to less favorable protein behavior during purification. Alternatively, therefore, we expressed Chl1 encoded by its native sequence as detailed below. Once purified, both Chl1 preparations showed similar behavior and activity, and they were used interchangeably in this study.

Budding yeast cells overexpressing Chl1 from its native sequences were grown in YP medium containing 2% raffinose as the carbon source to an optical density of 1.0 at 30 °C. 2% galactose was added to the culture to induce protein expression, and cells were further grown for 3 hours. Cells were collected by centrifugation, washed with deionized water, and suspended in Chl1 buffer (20 mM HEPES-KOH pH 7.5, 20% glycerol, 300 mM NaCl, 0.01% Tween-20, 0.5 mM TCEP) containing 0.5 mM Pefabloc, as well as SIGMAFAST Protease Inhibitor Cocktail EDTA-free. The cell suspension was frozen in liquid nitrogen, then cells were broken in a cryogenic freezer mill. The cell powder was thawed on ice, and further Chl1 buffer containing protease inhibitors was added. The lysate was clarified by centrifugation at 20,000 x g for 1 hour. The lysate was transferred to pre-equilibrated anti-FLAG M2 affinity gel and incubated for 2 hours. The resin was washed with Chl1 buffer, then incubated in the same buffer containing 10 mM MgCl_2_ and 1 mM ATP for 15 minutes. The resin was washed again with Chl1 buffer, then incubated in the same buffer containing 0.5 mg/ml FLAG peptide. The eluate was loaded onto a Superdex 200 Increase 10/300 GL gel filtration column that was equilibrated and developed with Chl1 gel filtration buffer (20 mM HEPES-KOH pH 7.5, 200 mM NaCl, 10% Glycerol, 0.5 mM TCEP). The peak fractions were pooled.

### DNA substrates

pBluescript-based DNA replication substrates, harboring the *ARS1* replication origin and a size of 3.2 kb or 5.8 kb ^33^ were prepared from *E. coli* using the QIAGEN Plasmid Mega Kit. While the resultant preparation majorly consists of monomeric, covalently closed circular DNA, a minority of nicked or catenated species are observed, which were removed as follows. To eliminate nicked DNA, the preparation was treated with T5 exonuclease, and the reaction quenched by addition of SDS to a final concentration of 1%. The DNA was then loaded onto a 10 - 25% sucrose gradient, in gradient buffer (50 mM Tris-HCl pH 8, 2 mM EDTA, 50 mM NaCl, 0.1% Triton X-100) prepared in 5 ml Open-Top Thinwall Ultra-Clear Tubes. The tubes were centrifuged at 73,000 x g for 18 hours (for the 3.2 kb plasmid) or at 58,000 x g for 16 hours (for the 5.8 kb plasmid). 0.3 ml fractions were then harvested manually. Monomeric DNA peak fractions were identified by agarose gel electrophoresis, and the DNA purified and recovered using phenol-chloroform extraction and ethanol precipitation.

### Cohesion establishment using pre-loaded cohesin

All incubations were conducted at 30 °C. Cohesin loading was performed in CL buffer (25 mM HEPES-KOH pH 7.5, 10% Glycerol, 0.5 mM MgCl_2_, 0.003% NP-40, 2 mM TCEP, 20 mM potassium glutamate, 5 mM ATP). Circular DNA, cohesin and the Scc2-Scc4 cohesin loader complex were added to final concentrations of 3.3 nM, 120 nM and 60 nM, respectively. The reaction proceeded for 30 minutes and then 1.5 times diluted in MCM loading buffer (25 mM HEPES-KOH pH 7.5, 30 mM magnesium acetate, 2 mM TCEP, 0.02% NP-40, 250 mM potassium glutamate, 10 mM ATP). ORC, Cdt1-Mcm2-7 and Cdc6 were added at a final concentration of 10 nM, 30 nM, 50 nM, respectively. After 15 minutes incubation, DDK and S-CDK were added at a final concentration of 40 nM and 20 nM, respectively. After further 15 minutes, the reaction was twofold diluted in replication buffer (25 mM HEPES-KOH pH 7.5, 300 mM potassium glutamate, 10 mM magnesium acetate, 0.02% NP-40, 2.4 mM ATP, 400 μM each of TTP, GTP and CTP, 80μM each of dATP, dTTP, dGTP and dCTP, 2 mM TCEP, 1 mM acetyl-CoA, 66 nM [α-^32^P]dCTP). If not indicated otherwise, 100 nM RPA, 30 nM Dpb11, 40 nM Cdc45, 5 nM GINS, 10 nM S-CDK, 10 nM Top I, 20 nM Ctf4, 40 nM Chl1, 20 nM Polε, 5 nM Mcm10, 20 nM Tof1-Csm3, 20 nM Mrc1, 40 nM RFC, 80 nM PCNA, 0.4 nM Pol δ, 40 nM Fen1, 60 nM Cdc9, 50 nM Pds5, 40 nM Eco1, 10 nM Dna2, 25 nM Pol α, 65 nM Sld2, 25 nM Sld3-Sld7, 10 nM Ctf18-RFC, 5 nM Pif1, and either 2.5 nM budding yeast Top II or 2 nM *Chlorella* virus Top II were added, and replication reactions were incubated for 40 minutes.

### Cohesin immunoprecipitation (without crosslinking)

8 μl of a replication reaction was taken as the DNA input sample. 12 μl deproteination buffer (10 mM Tris-HCl pH 7.5, 1 mM EDTA, 50 mM NaCl, 0.75% SDS) containing 1.7 mg/ml proteinase K were added, and incubated at 37 °C for 20 minutes. 80 μl replication reaction were used to immunoprecipitate cohesin. 250 μl of IP buffer 1 (25 mM HEPES-KOH pH 7.5, 500 mM NaCl, 0.1% NP-40, 5 mM EDTA) were added together with α-Pk antibody-coated protein A-conjugated magnetic beads. The mix was incubated on ice with gentle agitation for 40 minutes, and then washed extensively with IP buffer 1. The magnetic beads were then suspended in 20 μl of deproteination buffer containing 1 mg/ml proteinase K, and incubated at 37 °C for 20 minutes.

The input and immunoprecipitated DNA fractions were phenol-chloroform extracted, then loaded onto MicroSpin G-50 Columns, equilibrated with Nb.BssSI buffer (50 mM Tris-HCl pH 8, 100 mM NaCl, 10 mM MgCl_2_, 0.02% NP-40). To nick DNA, if applicable, Nb.BssSI was added to and incubated for 20 minutes at 37 °C. The DNA was resolved by 0.8% agarose gel electrophoresis in TAE buffer and visualized using SYBR Gold Nucleic Acid Gel Stain and a ChemiDoc MP Imaging system (BioRad). To visualize replication products that incorporated [α-^32^P]dCTP, the same agarose gel was dried, exposed to an Imaging Plate (Fujifilm), and scanned using a Typhoon FLA 9500 biomolecular imager (Cytiva).

### 6C Cohesin crosslinking, denaturation and immunoprecipitation

DNA replication was conducted essentially as described above, but [α-^32^P]dCTP was used at a final concentration of 250 nM, and replication reaction proceeded for 50 minutes. Cdc9 was not included in the experiments shown in Figures 2B, 3D and 4B (to yield nicked replication products). After replication, 0.5 mM bis-maleimidoethane BMOE was added to 6C (or 5C) cohesin containing reactions. Crosslinking proceeded for 20 minutes at room temperature. The reaction was quenched by addition of 2 mM DTT. For protein denaturation, SDS was added to a final concentration of 1% before incubation at 65 °C for 20 minutes. The SDS-containing samples were diluted by tenfold using IP dilution buffer (25 mM Tris pH8, 150 mM NaCl, 1% NP-40, 20 mM EDTA), and BSA was added to a final concentration of 0.5 mg/ml. α-Pk antibody-coated protein A-conjugated magnetic beads were added, and the samples kept on ice with gentle agitation for 60 minutes. The magnetic beads were washed using wash buffer 1 (50mM Tris-HCl pH8, 500 mM NaCl, 10 mM EDTA, 0.1% NP-40). As applicable, to nick recovered DNA, the cohesin-DNA complexes on magnetic beads were further treated with Nb.BssSI buffer containing 200 U/ml Nb.BssSI for 15 minutes at 30 °C. To analyze the DNA methylation status, the cohesin-DNA complexes on magnetic beads were further treated in Dpn I buffer (50 mM Tris-HCl pH 8, 300 mM NaCl, 10 mM MgCl_2_, 0.05% NP-40) containing 200 U/ml Dpn I for 15 minutes at 30 °C. Cohesin-bound replication products were then eluted from the antibody beads in buffer containing 10 mM Tris-HCl pH 8, 1 mM EDTA, 100 mM NaCl, and 1% SDS at 65 °C.

### Two-dimensional (2D) agarose gel electrophoresis

6C cohesin-replicated DNA complexes were firstly resolved at 1.4V/cm on a 0.7% agarose TAE gel containing 0.2% SDS for 15 hours (first dimension). Lanes from the gel were cut and placed at the top of a second 0.7% agarose TAE gel containing 0.2% SDS, leaving an approximately 1.5 cm wide gap between the gel lane and the new gel. The gap was filled with 60 °C 0.7% agarose in TAE, containing 0.2% SDS and 0.2 mg/ml proteinase K. Once the molten agarose solution solidified, the 6C cohesin-replicated DNA complexes were again resolved at 1.4V/cm for 15 hours, where 6C cohesin was digested by proteinase K (second dimension). 0.2% SDS was included in the TAE running buffer for both first and second dimension. The agarose gel was dried, exposed to an imaging plate and scanned as above.

### Spontaneous cohesin loading without the cohesin loader

50 nM cohesin was incubated with 3.3 nM circular DNA in CL buffer for 60 minutes at 30 °C. The cohesin loading reaction was terminated by adding equal volume of IP buffer 2 (35 mM Tris-HCl pH 7.5, 180 mM NaCl, 20 mM EDTA, 10% glycerol, 0.05% NP-40). anti-FLAG M2 antibody-coated protein G-conjugated magnetic beads were added to the reaction, then the samples were rocked for 60 minutes at 4 °C. The magnetic beads were washed in wash buffer 2 (25 mM HEPES-KOH pH 7.5, 500mM NaCl, 0.02% NP-40, 5 mM EDTA, 1 mM TCEP), and then in MCM loading buffer (25 mM HEPES-KOH pH 7.5, 10 mM magnesium acetate, 0.01% NP-40, 100 mM potassium glutamate) containing 2 mM TCEP. The beads were suspended in the same buffer containing 0.1 mg/ml FLAG peptide, and incubated at 30 °C for 5 minutes. The beads were removed using a magnet, ATP was added to a final concentration of 5 mM, and MCM loading, DNA replication, cohesin crosslinking, protein denaturation, and cohesin immunoprecipitation were performed as above.

### Replication-dependent cohesin loading

4 nM 3.5 kb template DNA, 10 nM ORC, 50 nM Cdc6 and 30 nM Cdt1-Mcm2-7 were mixed in MCM loading buffer containing 0.5 mM TCEP and 5 mM ATP and incubated at 30 °C for 15 minutes. DDK and S-CDK were then added to a final concentration of 50 nM and 20 nM, respectively. After 15 minutes incubation, the reaction was twofold diluted into low-salt replication buffer (25 mM HEPES-KOH pH 7.5, 100 mM potassium glutamate, 10 mM magnesium acetate, 0.02% NP-40, 2.4 mM ATP, 400μM each of TTP, GTP and CTP, 80 μM each of dATP, dTTP, dGTP and dCTP, 2 mM TCEP, 1 mM acetyl-CoA, 66 nM [α-^32^P]dCTP). 100 nM RPA, 30 nM Dpb11, 40 nM Cdc45, 5 nM GINS, 10 nM S-CDK, 20 nM Top I, 20 nM Ctf4, 40 nM Chl1, 20 nM Polε, 5 nM Mcm10, 20 nM Tof1-Csm3, 20 nM Mrc1, 40 nM RFC, 80 nM PCNA, 25 nM Pol α, 65 nM Sld2, 25 nM Sld3-Sld7 were also added, together with 50 nM Cohesin and 30 nM Scc2-Scc4. The replication reaction proceeded for 20 minutes. (5 nM Pif1 was included and replication reactions extended to 40 minutes to facilitate quantitation of completely replicated DNA circles in the experiments shown in Figure S6). After replication, a 7 μl aliquot was taken as the DNA input sample. 1% SDS and 2 mg/ml proteinase K final concentrations were added and the aliquots incubated at 37 °C for 20 minutes. To analyze cohesin-bound DNA, 170 μl of IP buffer 1 were added to 28 μl of the remaining replication reactions, together with α-Pk antibody-coated protein A-conjugated magnetic beads. Immunoprecipitation was conducted for 40 minutes. The magnetic beads were washed with the IP buffer 1, then suspended in MCM loading buffer containing 1% SDS and 2 mg/ml proteinase K at 37 °C for 20 minutes.

The input and cohesin-bound DNA fractions were phenol-chloroform extracted and analyzed by agarose gel electrophoresis in TAE buffer. SYBR-Gold staining and autoradiography were performed as above.

### Replication-coupled cohesion establishment

To analyze the establishment of sister chromatid cohesion by replication-coupled cohesin loading, 6C-cohesin was used and 0.4 nM Pol δ, 40 nM Fen1, 60 nM Cdc9, 50 nM Pds5, 40 nM Eco1, 10 nM Dna2, 10 nM Pif1, 10 nM Ctf18-RFC, and 1.25 nM *Chlorella* virus Top II were supplemented to the above replication-coupled cohesin loading reactions. The reactions were incubated for 50 minutes. BMOE crosslinking, protein denaturation, cohesin immunoprecipitation, Nb.BssSI nicking enzyme treatment, Dpn I treatment, and 2D agarose gel electrophoresis were then performed as described above.

## QUANTIFICATION AND STATISTICAL ANALYSIS

Conclusions in this study that are based on qualitative comparisons were all confirmed by experimental repeats. In these cases, representative experiments are shown. In cases where a quantitative comparison was warranted, three independent repeat experiments were performed. Cohesin-associated replication products were detected and quantified by autoradiography using Typhoon FLA 9500 biomolecular imager (Cytiva). Individual results from all three repeats are shown, together with the means and standard deviations.

---

**Figure S1.**
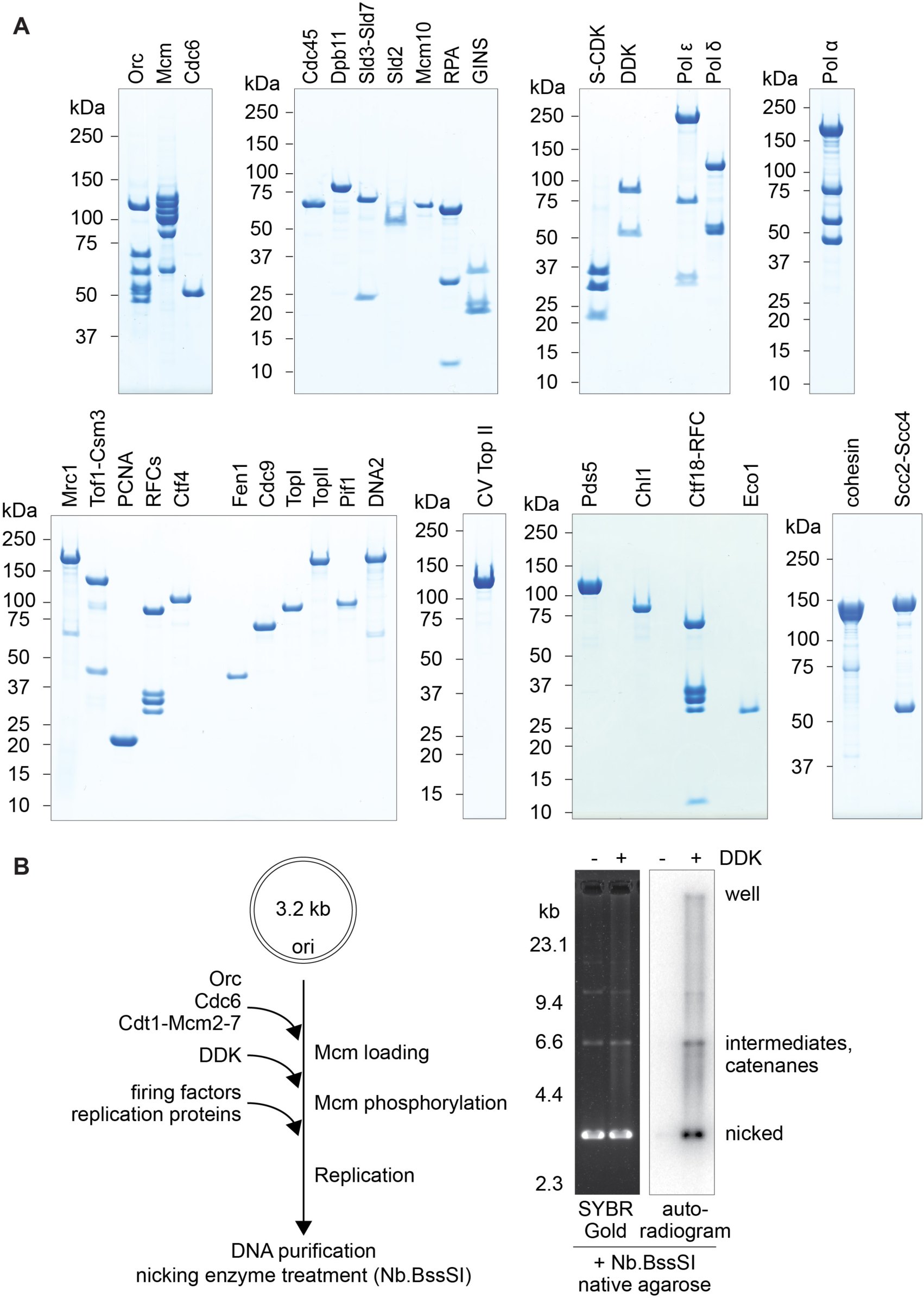
Characterization of the DNA replication reaction, Related to Figure 1. (A) Purified replication and sister chromatid cohesion factors used in this study were analyzed by SDS-PAGE followed by Coomassie Blue staining. (B) Schematic of a complete plasmid DNA replication reaction, and example gel electrophoretic analysis of the total DNA visualized with the DNA stain SYBR Gold and of the replication products visualized by [α-^32^P]dCTP autoradiography. While our input template DNA is supercoiled, topoisomerases present in the replication reaction lead to its partial relaxation. To facilitate the comparison between input and replicated DNA, irrespective of their supercoiling state, we treated all samples with a nicking enzyme before agarose gel electrophoresis that converts them into relaxed circles. The experiment demonstrates the dependence of DNA replication on the DDK origin activation kinase.

**Figure S2.**
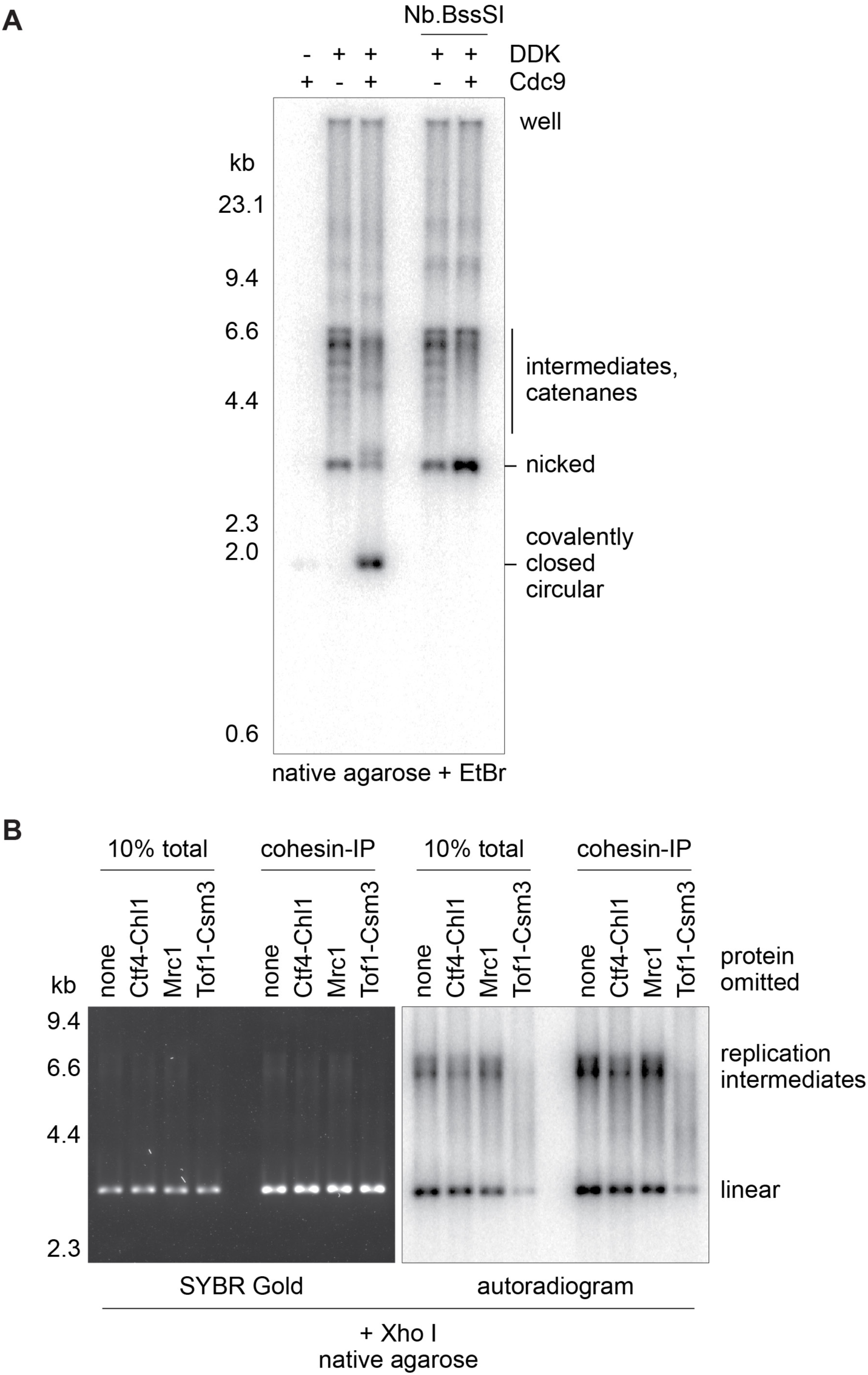
Characterization of the fully replicated species, and an experiment to analyze the contribution of cohesion establishment factors to cohesin retention during DNA replication, Related to Figure 1. (A) Evidence that the fast-migrating population of replication products, detected during agarose gel electrophoresis in the presence of ethidium bromide (EtBr), consists of fully replicated, covalently closed circular DNAs. The species does not form in the absence of the DNA ligase Cdc9, required to seal nicks that remain between processed Okazaki fragments (left). Furthermore, nicking of the product by Nb.BssSI treatment following DNA purification prevents supercoiling by EtBr intercalation. (B) The indicated cohesion establishment factors were omitted from the DNA replication reactions. Total DNA following DNA replication, as well as cohesin-bound fractions were analyzed. Replication products were visualized by [α-^32^P]dCTP autoradiography. To facilitate comparisons, DNAs were cleaved with the restriction enzyme XhoI before gel electrophoresis, resulting in a single band representing fully replicated products. Note that, as expected^34^, the efficiency of DNA replication was markedly reduced in the absence of Tof1-Csm3.

**Figure S3.**
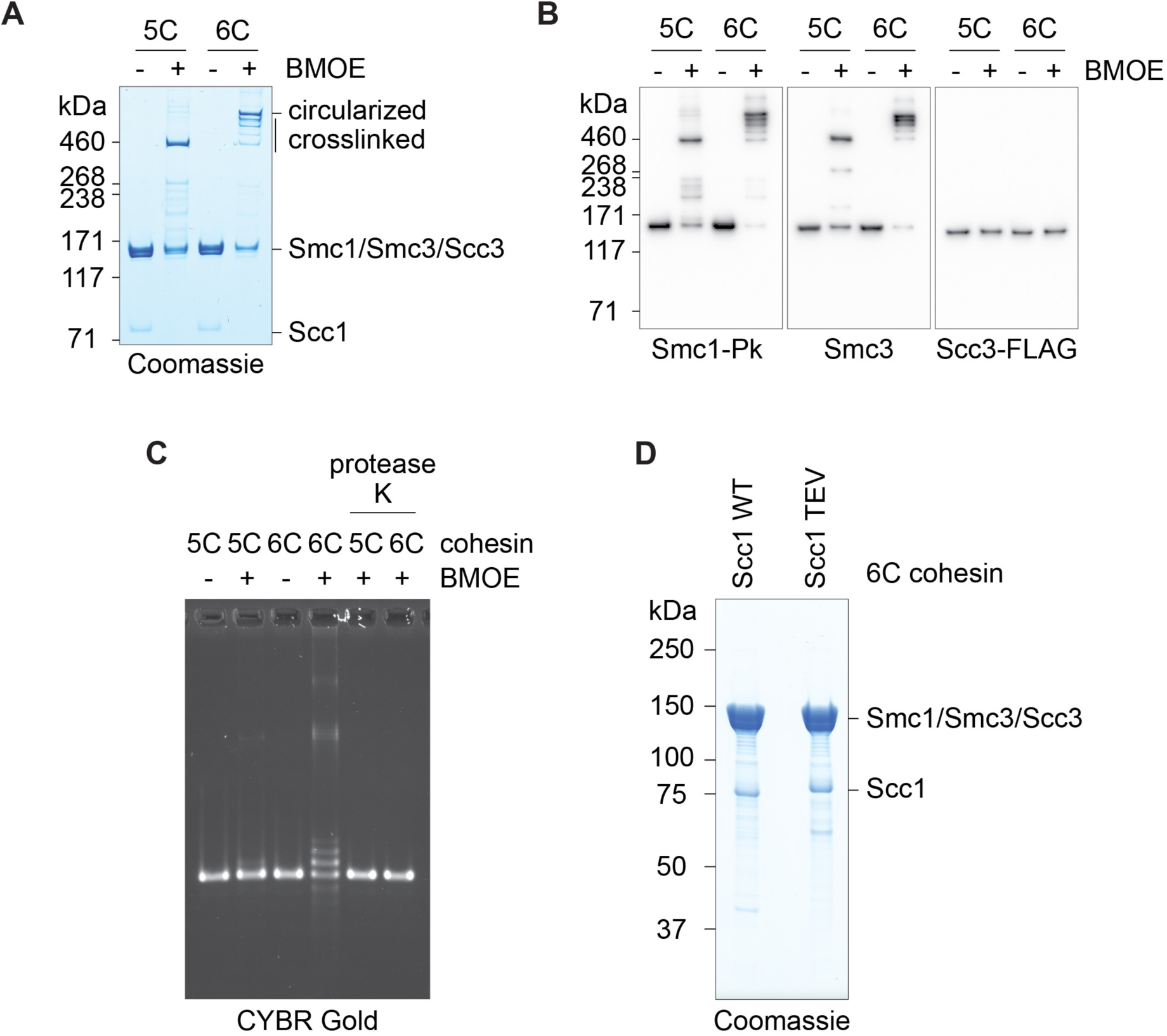
Characterization of 6C cohesin crosslinking, Related to Figure 2. (A) 5C and 6C cohesin were incubated in the absence or presence of bis-maleimidoethane (BMOE) and analyzed by SDS-PAGE followed by Coomassie staining. (B) Identification of the crosslinked species. As (A), but immunoblotting was used to detect the indicated subunits using their respective antibodies. (C) Topological DNA entrapment by 6C cohesin. 5C and 6C cohesin were loaded onto the circular template DNA in presence of the cohesin loader, followed by BMOE crosslinking as indicated, and SDS denaturation. Where indicated proteinase K was included before the DNA was resolved by agarose gel electrophoresis. (D) Purified 6C cohesin containing wild type Scc1 (WT), or Scc1 in which one of the separase recognition sites was exchanged for the TEV protease recognition sequence (TEV)^38^ were analyzed by SDS-PAGE followed by Coomassie staining.

**Figure S4.**
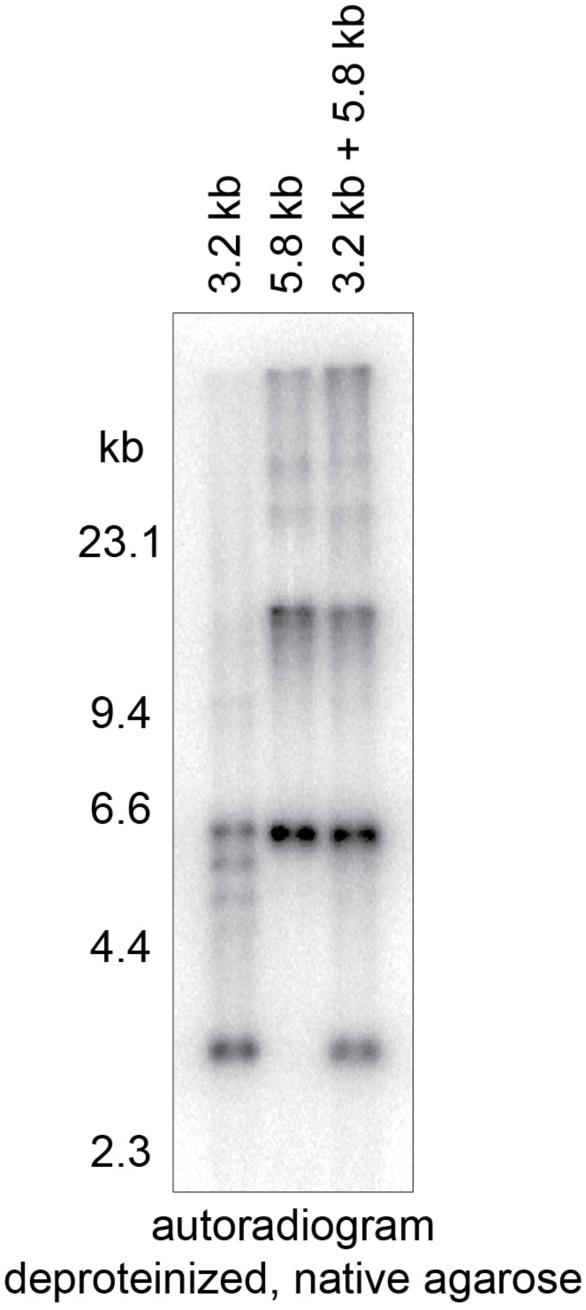
*In vitro* replication of two DNA templates of different sizes, Related to Figure 3. Replication reactions were performed with either or both of two circular, *ARS1*-containing, DNA templates 3.2 kb and 5.8 kb in size. Replication products were analyzed following deproteinization by agarose gel electrophoresis and autoradiography. When combined, both templates were replicated with approximately equal efficiencies.

**Figure S5.**
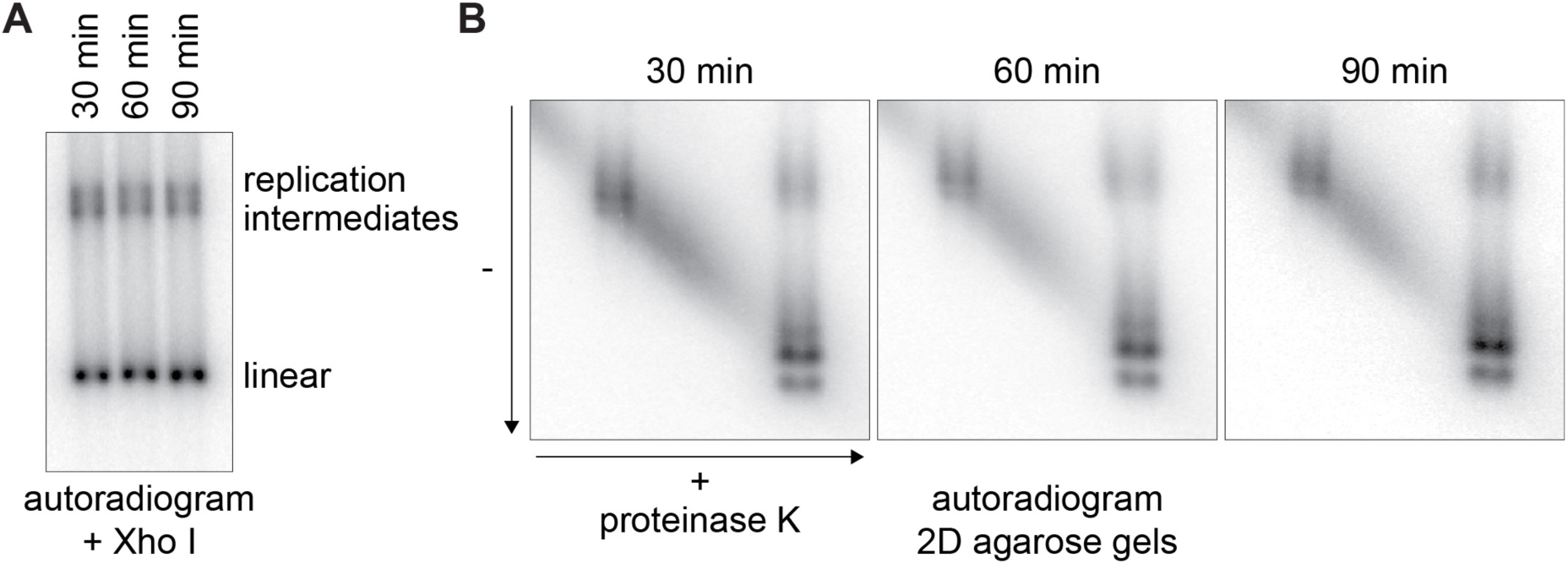
Timecourse analysis to probe the stability of sister chromatid cohesion established during *in vitro* DNA replication, Related to Figure 4. (A) Replication reactions using 6C cohesin-bound templates were incubated for the indicated times. The replication products were analyzed following deproteinization and restriction enzyme cleavage using XhoI. This analysis reveals that replication has largely reached completion after 30 minutes. (B) Aliquots from the replication reactions in (A) were BMOE crosslinked, denatured, and cohesin-bound DNAs were enriched by cohesin immunoprecipitation. 2D gel electrophoresis reveals that the fraction of cohesin that promotes sister chromatid cohesion, or associates with only one of two replication products, remains constant over time.

**Figure S6.**
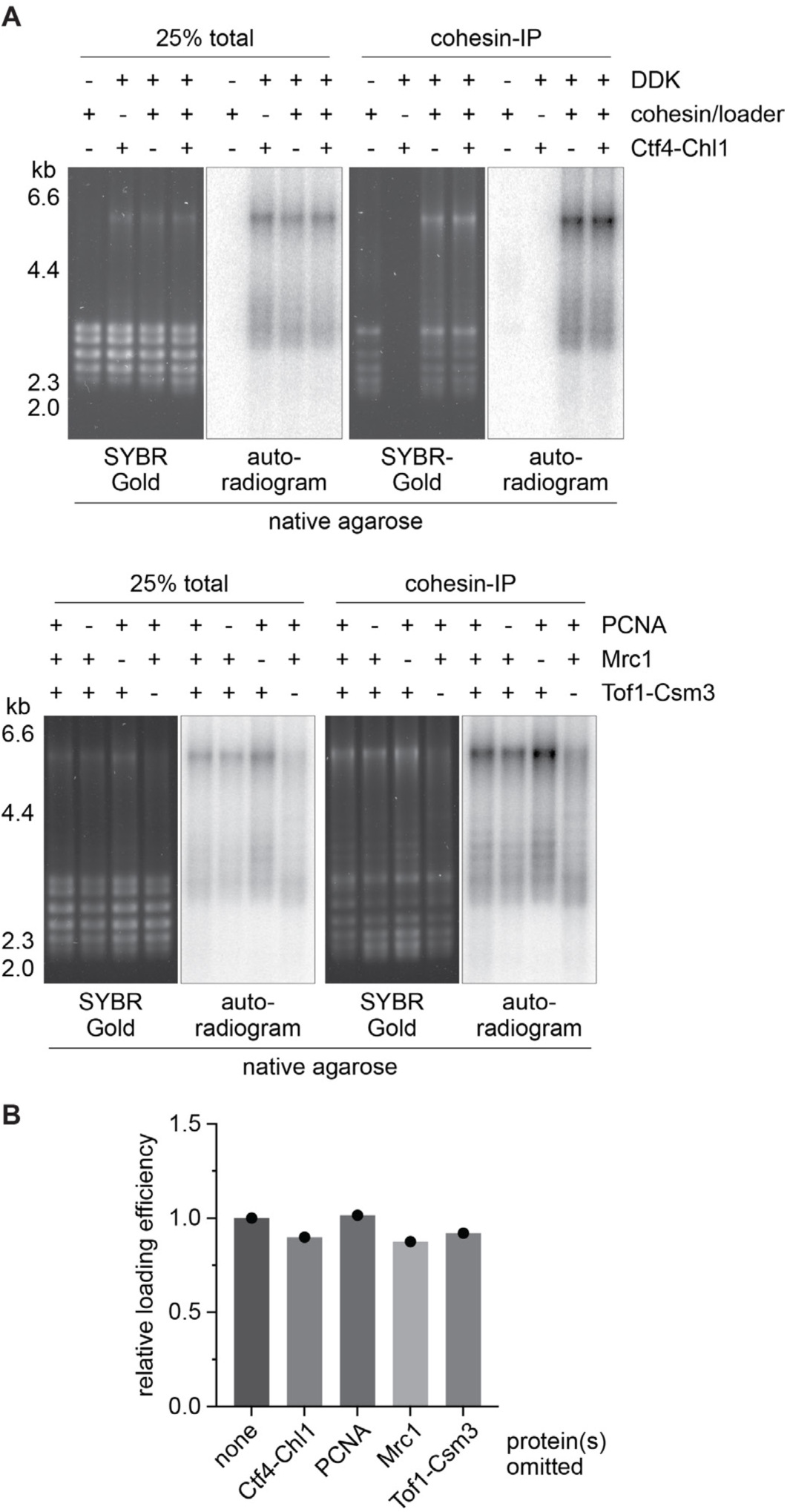
Characterization of replication-coupled cohesin loading, Related to Figure 5. (A) Replication reactions were performed in the presence of cohesin with or without the indicated factors. Total and cohesin-bound DNAs were afterwards analyzed by agarose gel electrophoresis. SYBR Gold stained all DNA, while autoradiography was used to detect replication products. (B) The replication-coupled cohesin loading efficiency was quantified as the fraction of replication products retrieved by cohesin immunoprecipitation and is shown relative to a reaction in which no protein was omitted.

